# B2LiVe, a label-free 1D-NMR method to quantify the binding of amphitropic peptides or proteins to membrane vesicles

**DOI:** 10.1101/2022.12.14.520471

**Authors:** Mirko Sadi, Nicolas Carvalho, Bruno Vitorge, Daniel Ladant, J. Iñaki Guijarro, Alexandre Chenal

## Abstract

Amphitropic proteins and peptides reversibly partition from solution to membrane, a key process that regulates their functions. Experimental approaches, such as fluorescence and circular dichroism, are classically used to measure the partitioning of amphitropic peptides and proteins into lipid bilayers, yet hardly usable when the peptides or proteins do not exhibit significant polarity and/or conformational changes upon membrane binding. Here, we describe B2LiVe (*i*.*e*., Binding to Lipid Vesicles), a simple, robust, and widely applicable NMR method to determine the solution-to-membrane partitioning of unlabeled proteins or peptides. The experimental strategy proposed here relies on previously described proton 1D NMR fast pulsing techniques with selective adiabatic pulses. Membrane partitioning induces a large line broadening leading to a progressive loss of protein signals, and therefore, the decrease of the NMR signal directly measures the fraction of membrane-bound protein. The B2LiVe method uses low polypeptide concentrations and has been validated on several membrane-interacting peptides and proteins, ranging from 3 to 54 kDa, with membrane vesicles of different sizes and various lipid compositions.

**Motivation:** Characterization of the interaction of peptides and proteins with lipid membranes is involved in various biological processes and is often challenging for polypeptides which do not possess intrinsic fluorophores, do not exhibit significant structural content changes, as well as for those characterized by low affinities for membranes. To meet these challenges, we have developed a simple and robust label-free NMR-based experimental approach, named B2LiVe, to measure the binding of polypeptides to lipid vesicles. The experimental strategy relies on previously described proton 1D NMR fast pulsing techniques with selective adiabatic pulses to excite the amide resonances. B2LiVe is a label-free method based on the observation of amide hydrogen nuclei which are naturally present in all protein and peptide backbones. Our results validate the B2LiVe method and indicate that it compares well with established technics to quantify polypeptide-membrane interactions. Overall, B2LiVe should efficiently complement the arsenal of label-free biophysical assays available to characterize protein-membrane interactions.

**In brief:** We describe a robust label-free NMR-based experimental approach (B2LiVe) to measure interactions between peptides or proteins with membranes. The validity of this approach has been established on several polypeptides and on various membrane vesicles. The B2LiVe method efficiently complements the arsenal of label-free biophysical techniques to characterize protein-membrane interactions.

**Highlights:** B2LiVe is a simple and robust NMR-based method to quantify affinity of proteins and peptides for membranes

B2LiVe is a label-free approach that relies on proton 1D NMR fast pulsing techniques with selective excitation of amide resonances

B2LiVe has been validated on several membrane-interacting peptides and proteins

B2LiVe coupled to DOSY can pinpoint the presence within a membrane-bound protein of polypeptide segments remaining in solution

## Introduction

Characterization of the interaction of peptides and proteins with lipid membranes often begins with the determination of the affinity, or more appropriately, the partition coefficient Kx. A common preliminary study consists in the identification of the lipid species as well as the lipid properties, such as lipid polymorphism, charge, and acyl chain fluidity, favoring the partitioning of proteins and peptides from solution to membrane. Moreover, knowledge of membrane affinity can be important to decipher molecular mechanisms or mutational analyses to identify regions or amino acid residues critical for membrane binding.

Several experimental approaches, such as SPR (Surface Plasmon Resonance), fluorescence, centrifugation and circular dichroism are commonly used to measure the partitioning of soluble peptides and proteins into lipid bilayers. However, some peptides and proteins do not possess intrinsic fluorophores, or their secondary or tertiary structural contents do not change significantly enough to be used as a probe of their partitioning into membranes. Alternatively, they may exhibit a propensity to aggregate, complicating, if not precluding, the use of SPR or centrifugation-based approaches. Besides, techniques based on phase separation (e.g., centrifugation or membrane flotation assays) are hardly applicable for peptides/proteins showing low affinity for membranes.

In this article, we describe B2LiVe (Binding to Lipid Vesicles), a simple and robust NMR-based experimental approach to determine the solution-to-membrane partition coefficient, *K*_*x*_, of unlabeled peptides and proteins. The experimental strategy relies on proton 1D NMR fast pulsing techniques with selective adiabatic pulses to excite the amide resonances (HET-SOFAST) developed by Schanda and co-workers ^1^. Membrane partitioning induces a large line broadening leading to a loss of peptide/protein NMR signals. The decrease of the NMR signal thus directly reports the fraction of membrane-bound peptide/protein. The validity of this approach has been established on several membrane-interacting peptides and proteins, ranging from 3 to 54 kDa, and on membrane vesicles of various sizes and different lipidic compositions. The method does not require any labeling and requires only low amounts of peptide/protein when using high-field spectrometers and sensitive probes. Our results show that the B2LiVe NMR-based method compares well with traditional fluorescence and CD-based techniques. Overall, B2LiVe should efficiently complement the arsenal of label-free biophysical techniques to characterize protein-membrane interactions.

## Material and Methods

### Reagents

The lipids 1-palmitoyl-2-oleoyl-*sn*-glycero-3-phosphocholine (POPC, reference 850457C), 1-palmitoyl-2-oleoyl-*sn*-glycero-3-[phospho-*rac*-(1-glycerol)] (POPG, reference 840457C) and cholesterol (Chol, reference 700000P) were purchased from Avanti Polar Lipids (Alabaster, AL, USA). HEPES-Na (reference H9897), NaCl (reference S5150) and CaCl_2_ (reference 21115) were purchased from Sigma Aldrich, USA. The D_2_O (reference D214) was purchased from Eurisotop, England.

### Peptides

The synthetic peptides were purchased as powder from Genosphere Biotech (France) and their purity (95 %) and composition were controlled by reverse-phase HPLC and MALDI-mass spectrometry, respectively. The peptides are capped on the N terminus with an acetyl group and on the C terminus with an amide group. The P233 and P414 peptides used in this study were derived from the CyaA protein ^2–4^. The P233 peptide ^5,6^, which corresponds to the H-helix of ACD (adenyl-cyclase domain), residues 233-254 of the CyaA toxin ^5,7^ has the following sequence: LDRERIDLLWKIARAGARSAVG. The P414 peptide is derived from the segment 414-440 of CyaA ^6^ and contains a F415W mutation (SWSLGEVSDMAAVEAAELEMTRQVLHA). Peptide powders were resuspended in 20 mM HEPES-Na, 150 mM NaCl, pH 7.4. Peptide concentrations were calculated from the absorbance at 280 nm subtracted by the absorbance at 320 nm of absorbance spectra recorded with a Jasco V630 spectrophotometer using Hellma cuvettes. Aliquots of the dissolved peptides were stored at −20°C.

### Proteins

Bovine serum albumin (BSA, reference A0281), bovine α-lactalbumin (BLA, reference L-6010) and apo-myoglobin (apo-Mb, reference A8673-5X1VL) were purchased as powders from Sigma-Aldrich, USA. BSA was resuspended in H_2_O, BLA was resuspended in 20 mM HEPES-Na, 20 mM NaCl, 2 mM EDTA, pH 7.5 and apo-Mb was resuspended in 20 mM HEPES-Na, 20 mM NaCl, pH 7.5. After resuspension, BSA, BLA and apo-Mb were filtered through 0.2 *μ*m syringe filters (reference F2504-8, Thermofisher Scientific). Anthrolysin O (ALO) production and purification were performed as described in ^8^. The Diphtheria toxin translocation domain (T) ^9^ was produced and purified as described in ^10,11^. The calcium-free apo-state of calmodulin, apo-CaM, was produced and purified as described elsewhere ^127^. The physical-chemical parameters of the proteins and peptides used in this study are reported in Tables S1 and S2, respectively.

### Lipid vesicle preparation

Multilamellar vesicles (MLVs), large unilamellar vesicles (LUVs) and small unilamellar vesicles (SUVs) were used in this study. Lipid vesicles were prepared by reverse phase evaporation as described elsewhere ^13–16^ at a lipid concentration of 40 mM. MLVs produced by reverse phase evaporation were submitted to extrusion through 1.2 *μ*m polycarbonate filters. LUVs were prepared by further extrusion through 0.4 and 0.2 *μ*m polycarbonate filters. The SUVs were obtained by sonication of filtered MLVs. The hydrodynamic diameters and dispersity were checked by dynamic light scattering (DLS) using a NanoZS instrument (Malvern Instruments, Orsay, France). Lipid vesicles were aliquoted and stored under argon at 4°C. Various lipid compositions were used: POPC:POPG at a 8:2 molar ratio; POPC:POPG at a 9:1 molar ratio, both in 20 mM HEPES-Na, 20 mM NaCl, pH 4; POPC:cholesterol at a 6:4 molar ratio in 20 mM HEPES-Na, 20 mM NaCl, pH 7; POPC:POPG:cholesterol at a 7:2:1 ratio in 20 mM HEPES-Na, 150 mM NaCl, pH 7.

### Sample preparation

The samples used for NMR, tryptophan fluorescence and far-UV CD were derived from a common stock solution to ensure the comparability of the three methods, when applicable. A typical titration series comprises 19 samples. Protein (3 *μ*M) and peptide (15 *μ*M) concentrations are kept constant for all samples, while the lipid concentration increases from 0 up to 10 mM. Apo-Mb and T titrations were performed at 25°C in the presence of POPC/POPG 8:2 (apo-Mb) or 9:1 LUV (T) in 20 mM HEPES-Na, 20 mM NaCl, 5% D_2_O, pH 4. The titrations of ALO were performed in the presence of POPC/cholesterol 6:4 LUVs in 20 mM HEPES-Na, 20 mM NaCl, 2 mM CaCl_2_, 5% D_2_O, pH 7.5 at 37°C. The titrations of the P233 and P414 peptides were performed at 25°C in the presence of POPC/POPG/cholesterol 7:2:1 SUVs in 20 mM HEPES-Na, 150 mM NaCl, 2 mM CaCl_2_, 5% D_2_O, pH 7.5.

### Nuclear Magnetic Resonance

NMR experiments were performed on an Avance Neo 800 MHz spectrometer (Bruker, Billerica, MA, USA) equipped with a triple resonance cryogenically cooled probe. Mono-dimensional (1D) proton experiments with selective excitation of the amide region ^17^ of peptide (5-15 *μ*M) and protein (1-3 *μ*M) samples in the presence of varying lipid concentrations were recorded at 25°C or 37°C as indicated. Experiments were based on the proton HET-SOFAST pulse sequence ^1^ implemented in the NMRLIB 2.0 package ^18^. Selective adiabatic polychromatic PC9 pulses with a 120° pulse centered at 9.5 ppm with a 3-4 ppm bandwidth were used to excite amide resonances. The recycling time between scans was 0.15 ms. Spectra were processed and analyzed with Topspin 4.0.7 (Bruker). The relative amount of peptide/protein remaining in solution at each lipid concentration was evaluated from the integral of the amide proton resonances, after subtraction of the spectrum of lipids at the highest concentration used, considered as the baseline. Integral errors were estimated from the spectral noise standard deviation summed over an equivalent region.

Self-diffusion experiments of the T domain (15 *μ*M) at pH 4.5 were performed in the presence or absence of 15 mM LUV (POPC:POPG 9:1) at 25°C by means of the amide selective self-diffusion experiment 1D_DOSY (Diffusion Ordered SpectroscopY) implemented in the NMRLIB 2.0 package ^18^. We used a diffusion delay of 140 ms, bipolar shaped gradients applied during 3.7 ms and 5 gradient strengths that varied from 2 to 98% of the probe maximum gradient (53.5 G/cm). The selective pulses were centered at 9.5 and had a 3.1-ppm width. Water was suppressed with an excitation sculpting scheme incorporated in the 1D_DOSY scheme. To evaluate the effect of the different viscosity of T samples ± LUV on diffusion, we followed the signal of Hepes at 3.89 ppm contained in the buffer in non-selective experiments. We recorded 16 stimulated-echo experiments with a diffusion delay of 60 ms, bipolar gradients applied during 1.6 ms at different gradient strength and excitation sculpting water suppression. Integral errors for each data point were estimated from the noise standard deviation.

### Intrinsic tryptophan fluorescence

Measurements were performed at 25 or 37 °C with a FP750 spectrofluorimeter (Jasco, Japan) equipped with a Peltier-thermostated cell holder, using a quartz cell (reference 105.251-QS, 3×3 mm pathlength) from Hellma (France). The peptide tryptophan emission spectra were recorded from 300 to 400 nm (excitation at 280 nm at a scan rate of 100 nm/min). A bandwidth of 5 nm was used for both, excitation and emission beams. The ratio of fluorescence intensities at 320 and 370 nm extracted from the fluorescence emission spectra was used to monitor solution to membrane partitioning.

### Determination of the partition coefficient Kx and of the free energy of partitioning ΔG_Kx_

The solution to membrane partitioning, *K*_*x*_, was determined by analysis of the fluorescence intensity ratios at 320/370 nm, of 1D NMR data and when applicable of CD data. The value of *K*_*x*_ is defined as the ratio of peptide/protein concentrations in membrane and solution phases ^19,20,14,6^, given by equation 1:

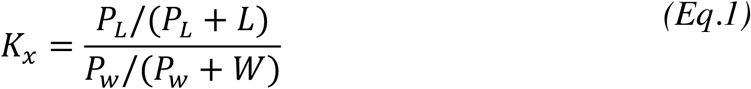

*P*_*w*_ (protein in aqueous phase) and *P*_*L*_ (protein in lipid phase) denote the concentration of soluble and membrane-bound protein, respectively; *P*_*T*_ stands for total protein (*P*_*T*_ = *P*_*w*_ + *P*_*L*_); *W* is the concentration of water (55.55 M); *L* is the concentration of lipid. Since *L* >> *P*_*L*_ and *W* >> *P*_*w*_, equation 1 can be written as follows:

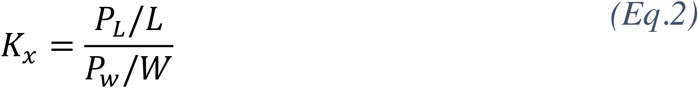

Equation 2 can, in turn, be expressed as follows:

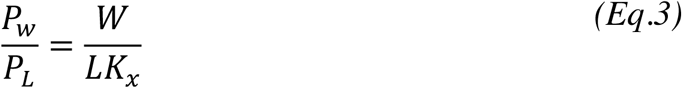

The peptide fraction f(P_L_) partitioned into membrane is described by the following equation:

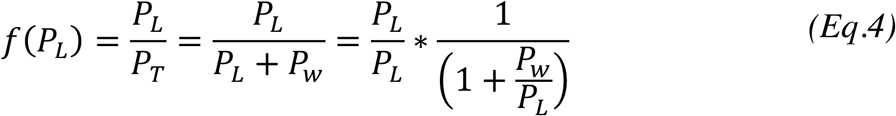

Based on equations 3 and 4, equation 5 can be expressed as follows:

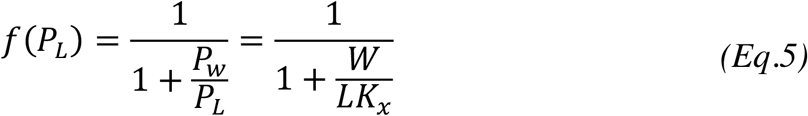

Adding the experimental data offset (A), the amplitude of the experimental signal (B) and the Hill coefficient (*n*) for the interaction to Equation 5, leads to Equation 6, which is used to fit all parameters to the experimental data:

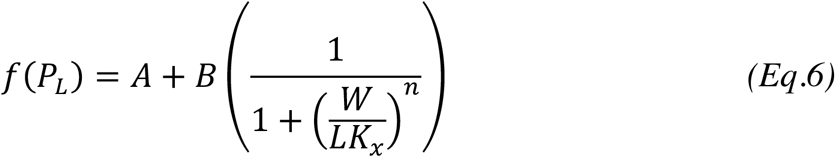

Equation 6 was fitted to the experimental data with Kaleidagraph (Synergy Software, Reading, USA).

The *K*_*x*_ constant can be expressed in terms of the dissociation constant *K*_*D*_ of the peptide-membrane interaction and the water concentration:

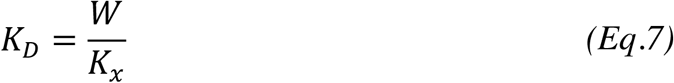

From Equation 7, Equation 6 rewrites:

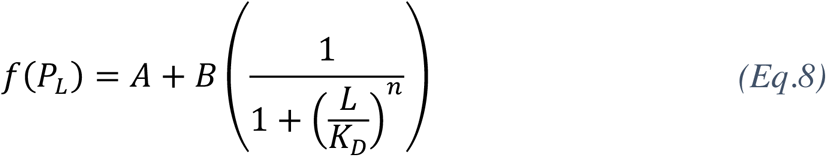

Finally, the free energy of solution to membrane partitioning Δ*G*_Kx_ was determined according to Equation 9:

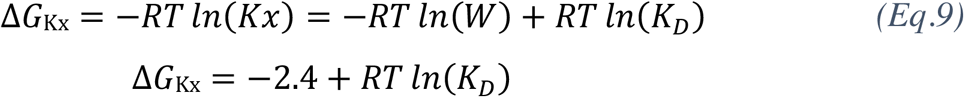

## Results

Protein binding to unilamellar vesicles of lipid bilayers induces a large line broadening in NMR signals, due to the slow tumbling rate of the particles, leading to a loss of protein signals. This observation has been exploited for characterization of the association of some ^15^N-labeled proteins with membranes ^21–23^. Here, we explored the possibility of quantifying membrane partitioning of unlabeled proteins by using proton 1D NMR fast pulsing techniques with selective excitation of amide protons using adiabatic pulses: the relative amount of protein or peptide remaining in solution at each lipid concentration should be proportional to the integral of the amide proton resonances. Hence, the decrease of the NMR signal should directly report on the fraction of protein bound to membranes.

To prove the general applicability of this approach, we investigated membrane partitioning of several proteins and peptides. Titrations were performed at constant protein and peptide concentrations and the lipid concentration was increased up to 10 mM lipids. Large and small unilamellar vesicles (LUV and SUV, respectively) of lipid bilayers were used. Protein partitioning into membranes was monitored by selective amide band 1D NMR (proton HET-SOFAST, see Methods), by recording the loss of intensity of the integral of the amide proton resonances. In parallel, protein binding to membranes was measured by intrinsic tryptophan fluorescence changes (following the ratio of fluorescence intensities emitted at 320 and 370 nm as a proxy for polarity change upon membrane binding), and far-UV circular dichroism (for peptides). The partition coefficient, *K*_*x*_, which is directly related to the affinity constant *K*_*D*_ (Equation 7), is determined by fitting Equation 8 to the experimental data (see Methods).

We first analyzed the partitioning of three amphitropic proteins into large unilamellar vesicles (LUV): apo-myoglobin (apo-Mb, 16.9 kDa) ^24,25^, the diphtheria toxin translocation domain (T, 22 kDa) ^26,27^, and anthrolysin O (ALO, 54 kDa) ^8^. Figure 1 shows the 1D NMR spectra of apo-Mb (Figure 1A), T (Figure 1B) and ALO (Figure 1C and 1D) with increasing concentrations of lipids.

**Figure 1.**
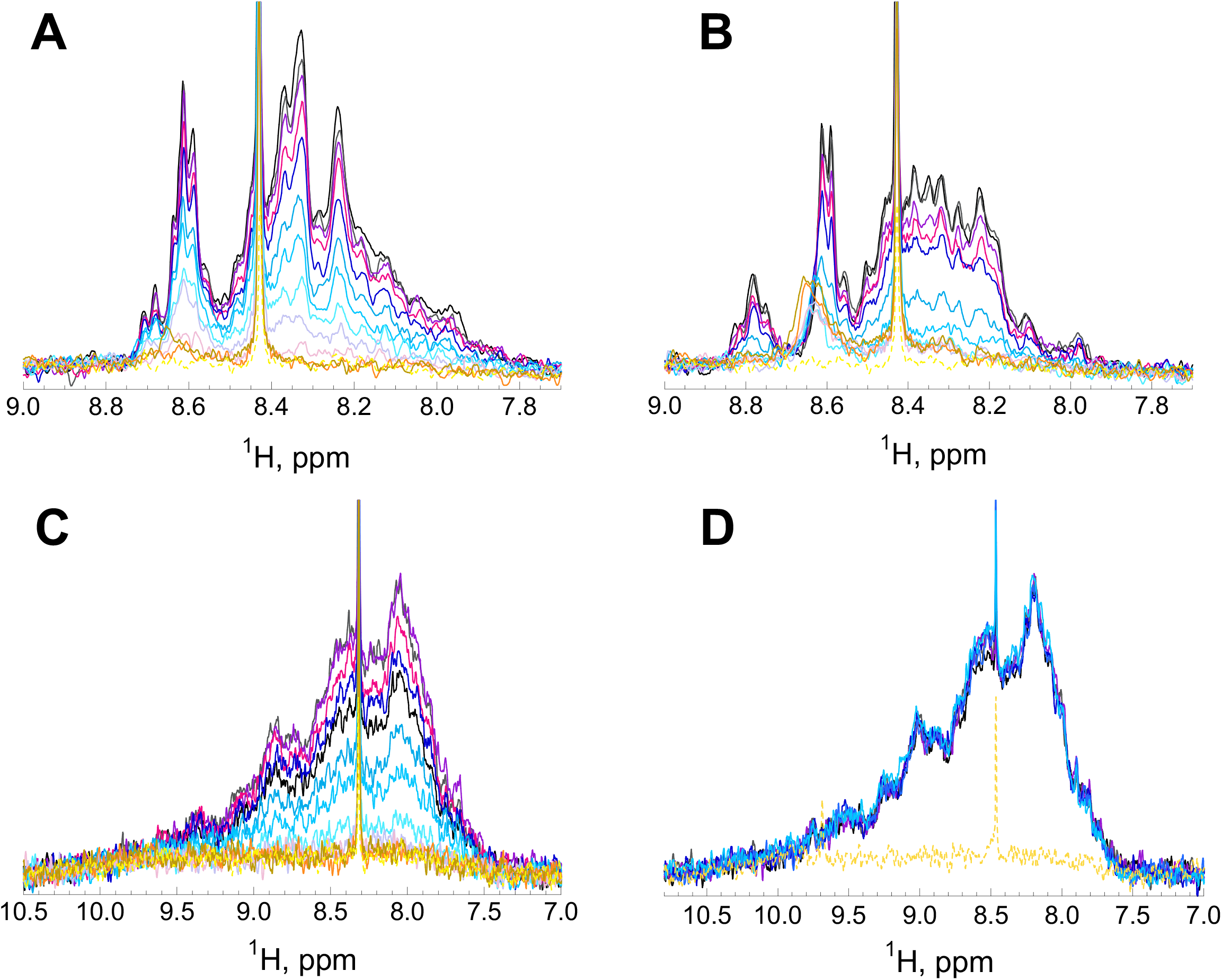
Membrane partitioning of amphitropic proteins monitored by 1D ^1^H amide-selective NMR spectroscopy. The concentration of lipids ranges from 0 mM (black) to 2 mM (dark yellow). The spectrum of lipid vesicles in the absence of protein at the highest lipid concentration used is indicated as a dashed line (yellow). Each protein is present at a concentration of 3 *μ*M. Membrane partitioning and corresponding NMR spectra upon titration of apo-Mb with POPC:POPG 9:1 LUV (A), T with POPC:POPG 9:1 LUV (B), ALO with POPC:cholesterol 6:4 LUV (C) and ALO with POPC LUV (D). Membrane partitioning is followed by the disappearance of the amide envelope signal (see main text for details). The sharp signal at 8.43 ppm arises from a trace impurity.

The intensity of the 1D NMR spectra of apo-Mb and T (Figure 1A-B) decreases with the addition of LUVs. ALO is a cholesterol-dependent cytolysin (CDC) that requires the presence of cholesterol in lipid bilayers to partition from solution to membranes. Whereas the 1D NMR spectra of ALO titrated by membranes composed of POPC and cholesterol show a loss of intensity (Figure 1C), the intensities of the 1D NMR spectra do not significantly change for ALO titrated by LUV composed of POPC only (Figure 1D). This clearly establishes that the loss of 1D NMR signal intensity is a straightforward indicator of ALO binding to membranes.

To estimate the partition coefficients of the different proteins, the fractions of lost intensity of the integral of the amide proton envelope of apo-Mb, T and ALO were plotted as a function of lipid concentrations as reported in Figure 2A. In parallel, membrane partitioning was monitored by recording the ratio of tryptophan fluorescence intensity for the same proteins as a function of lipid concentration (Figure 2B). The fluorescence data indicate that the three amphitropic proteins interact with membranes, as expected from the literature ^8,10,25^. Furthermore, fitting Equation 8 to 1D NMR and fluorescence experimental data provided similar partition coefficients for each protein (see Table S3). Hence, as NMR and fluorescence provide similar quantitative results, we conclude that 1D ^1^H NMR can reliably be used to report protein partitioning into membranes.

**Figure 2.**
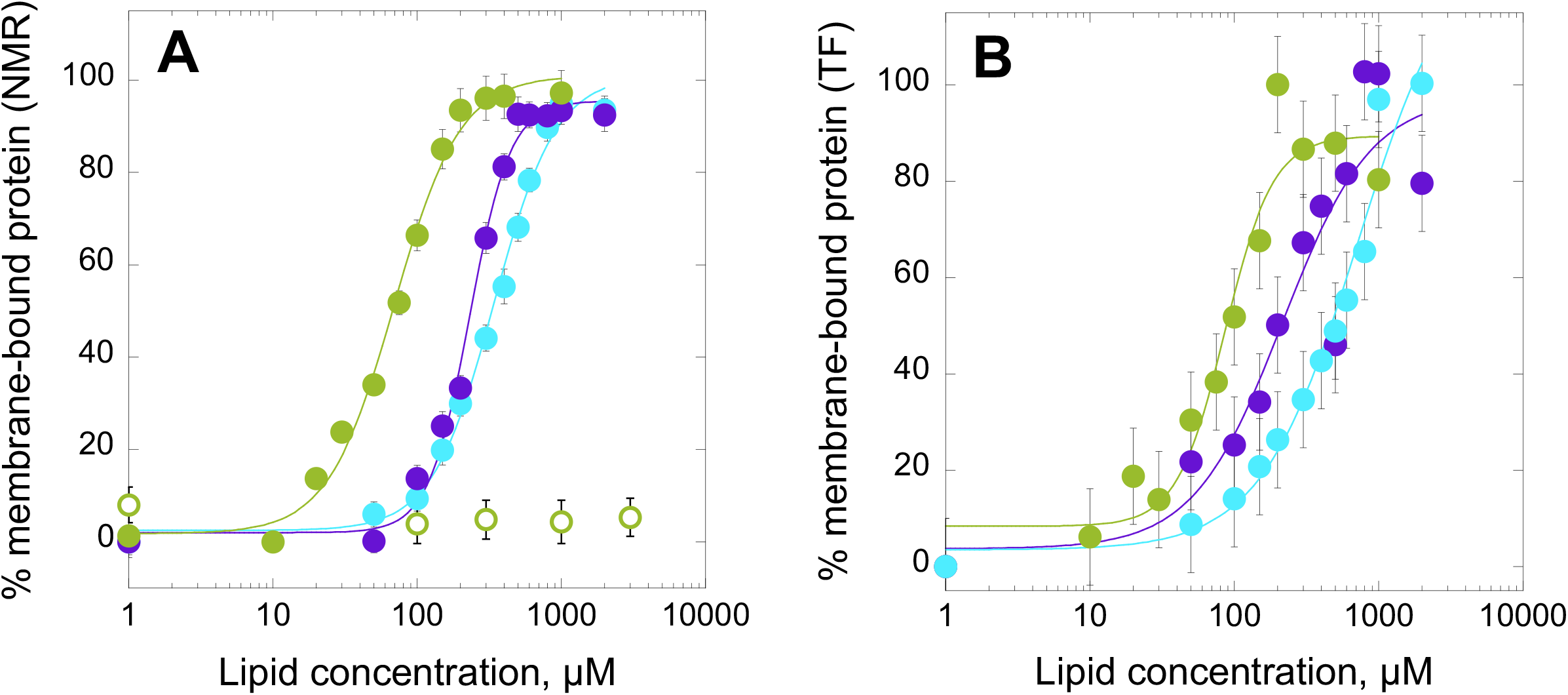
Membrane partitioning of amphitropic proteins. Solution-to-membrane partitioning of apo-Mb (cyan), T (violet) and ALO (green) at a concentration of 3 *μ*M, each in the presence of increasing lipid concentrations monitored by (A) 1D ^1^H NMR (signal loss of the amide envelope) and (B) fluorescence (ratio of tryptophan fluorescence intensities at 320 nm and 370 nm). POPC:POPG 9:1 LUV were used for apo-Mb and T, POPC:Chol 6:4 LUV for ALO (green, closed circles) and 100% POPC LUV for ALO (green, open circles). The error bars in (A) are calculated from the integral of the amide envelope (8.4-7.7 ppm) and from the spectral noise standard deviation, respectively.

As additional controls, we similarly characterized proteins that are not expected to interact with membranes of various lipid compositions: calmodulin (apo-CaM, 16.7 kDa), bovine holo-alpha-lactalbumin (hBLA, 14.2 kDa) and bovine serum albumin (BSA, 66.5 kDa) (Figure 3). These proteins were chosen to cover a similar range of molecular masses than apo-Mb, T and ALO. The addition of LUV does not affect the NMR signal nor the tryptophan fluorescence (the fluorescence experiment could not be done for CaM as it does not contain tryptophan), indicating that, as expected, these proteins do not interact with membranes under the experimental conditions used ^28^.

**Figure 3.**
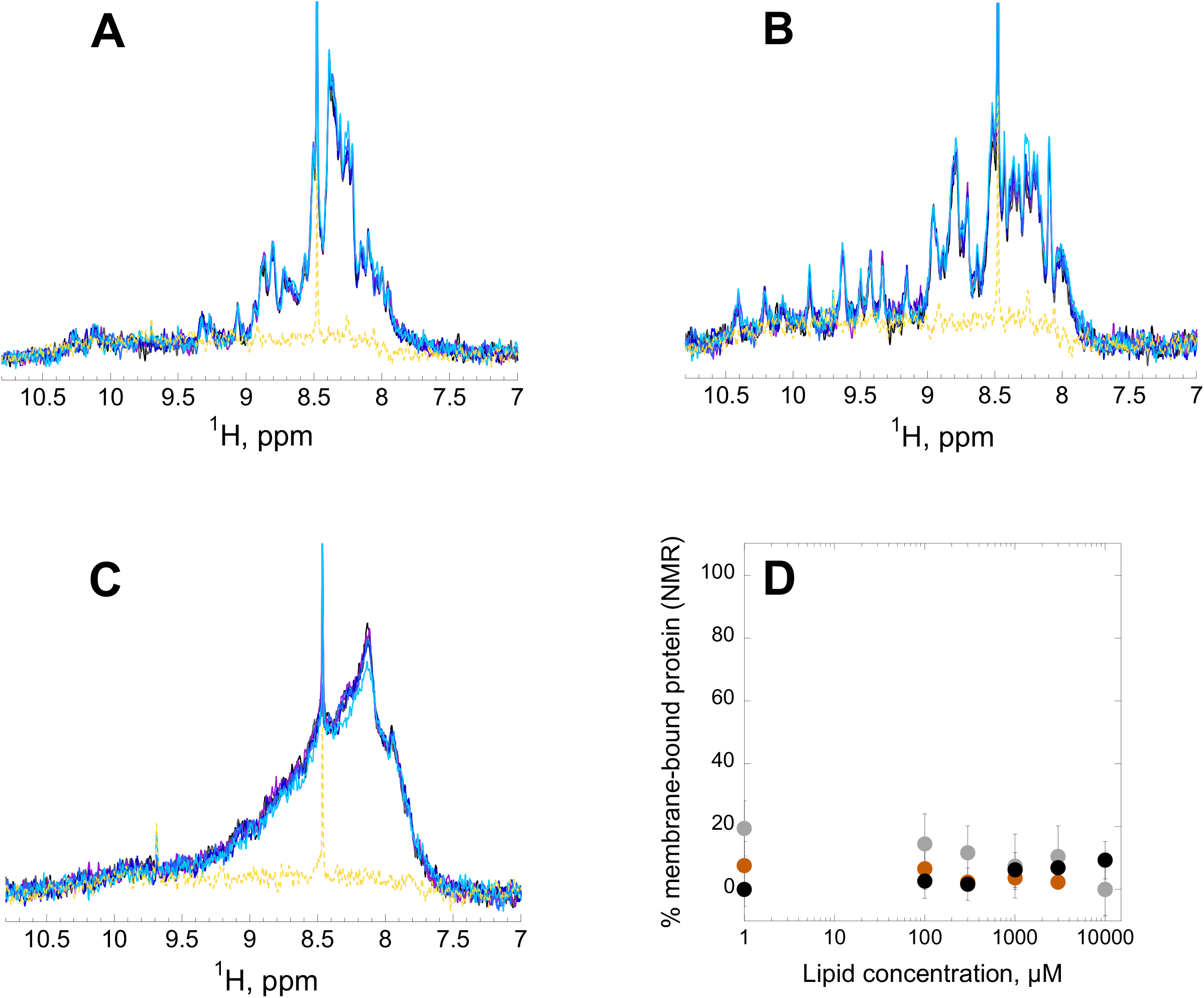
Selective amide spectra of soluble proteins that do not partition into membranes monitored by 1D ^1^H NMR spectroscopy. The concentration of lipids ranges from 0 mM (black) to 10 mM (light blue), the spectrum of lipid vesicles in the absence of protein at the highest lipid concentration used is indicated as a dashed line (yellow). Each protein is present at a concentration of 3 *μ*M. 1D ^1^H NMR spectra of proteins at increasing lipid concentrations are shown for apo-CaM in the presence of POPC:POPG 9:1 LUV (A), holo-BLA in the presence of POPC:POPG 9:1 LUV (B) and BSA in the presence of POPC:cholesterol 6:4 LUV (C). The sharp signals at 8.43 ppm in (A), (B) and (C) arise from an impurity. The plot in (D) represents the fraction of membrane-bound apo-CaM (brown), holo-BLA (grey) and BSA (black) in the presence of increasing lipid concentration. The fraction of membrane-bound proteins and the errors in (D) are calculated from the integral of the amide envelope (8.4-7.7 ppm) and from the spectral noise standard deviation, respectively.

Taken together, these results indicate that proton 1D NMR fast pulsing techniques with selective adiabatic pulses is a sensitive approach to quantitatively monitor protein partitioning into membranes. The decrease of the NMR signal directly reports on the fraction of membrane-bound proteins and can be easily recorded for unlabeled samples.

We then extended the study to peptide:membrane interactions. For this, we selected two peptides from the *Bordetella pertussis* adenylate cyclase toxin (CyaA)^3,4^, which were previously characterized in our lab: P233 that was shown to interact with membranes and P414 that has no membrane binding activity ^6^.

The NMR data shown in Figure 4 clearly indicate that P233 strongly interacts with POPC/POPG/Cholesterol membranes with a complete loss of NMR signal at high lipid/peptide ratios, while in the same conditions the P414 peptide does not show any signal decrease and therefore does not bind to membranes. Membrane partitioning of P233 was monitored in parallel by tryptophan fluorescence and far-UV circular dichroism (Figure 5). Overall, the experimental data revealed an excellent correspondence between the different techniques that provide similar quantitative parameters for the peptide membrane partitioning process (see Table S4). Hence, our data indicate that 1D ^1^H NMR is also readily applicable to quantify membrane partitioning of standard, unlabeled peptides.

**Figure 4.**
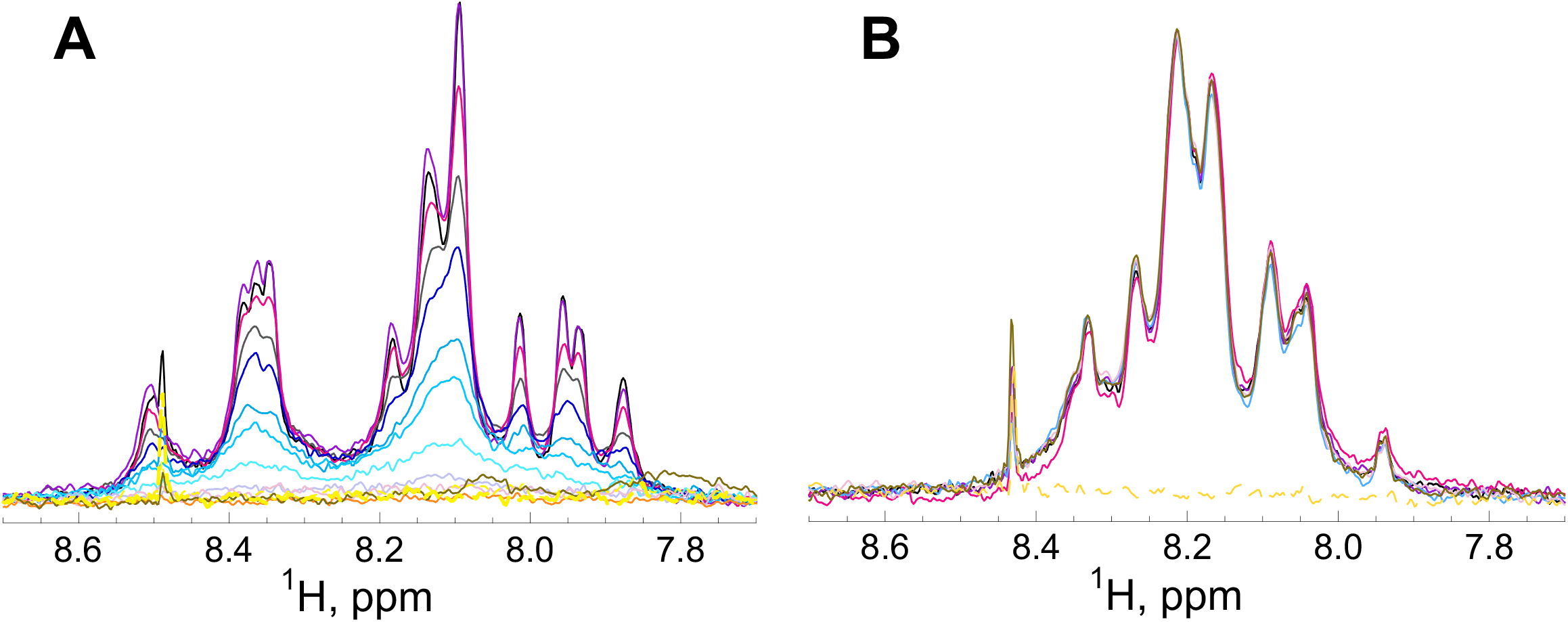
Membrane interaction of peptides followed by 1D ^1^H NMR spectroscopy. NMR amide spectra upon titration of the P233 (A) and P414 (B) peptides with POPC:POPG:cholesterol 7:2:1 SUV. The lipid concentrations range from 0 (black) to 10 mM (dark yellow). The concentrations of the P233 and P414 peptides are 15 *μ*M and 5 *μ*M, respectively. The spectrum of lipid vesicles in the absence of peptides at the highest lipid concentration used is represented as a dashed line (yellow).

**Figure 5.**
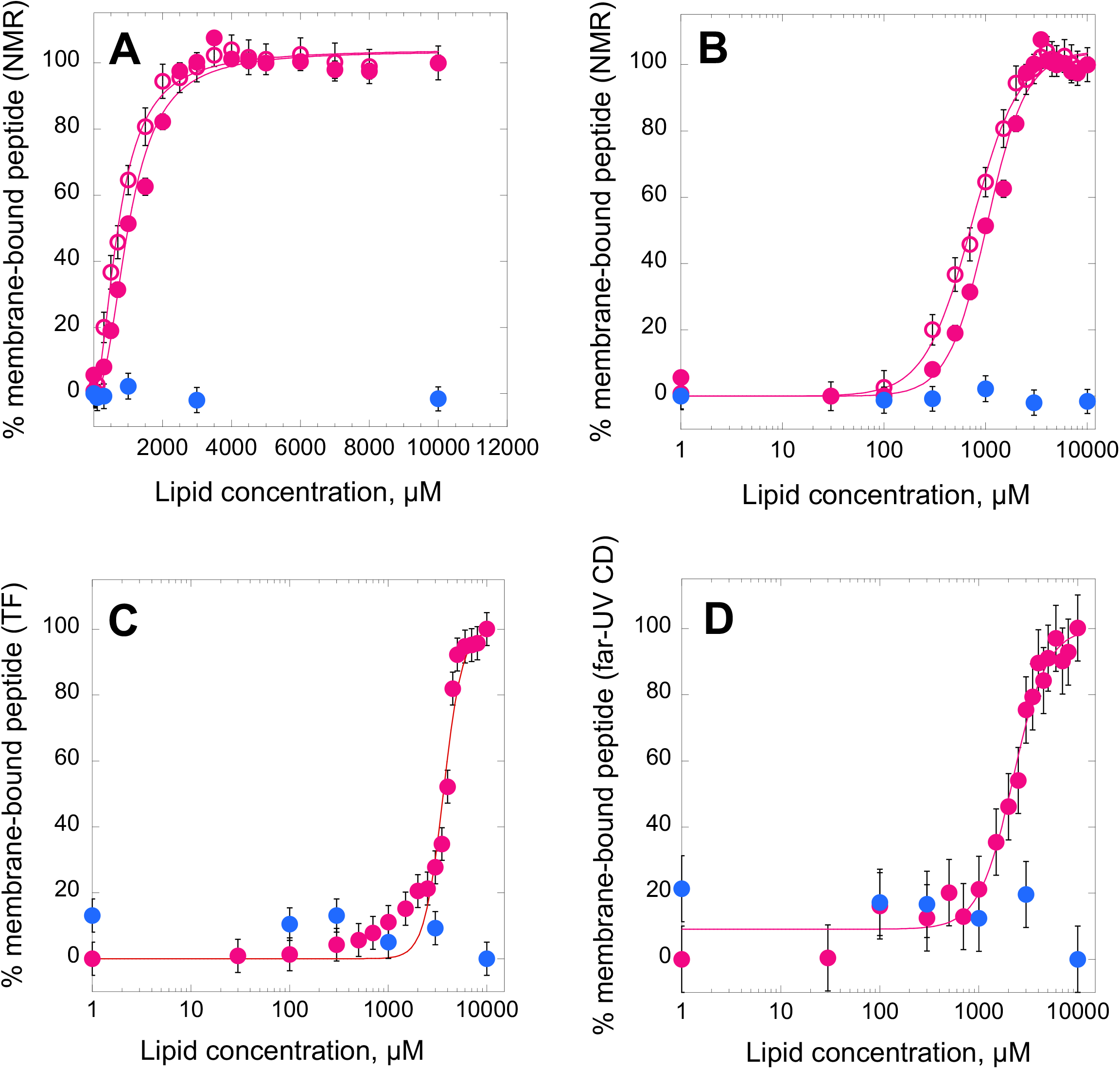
Membrane partitioning of the P233 and P414 peptides followed by 1D ^1^H NMR, tryptophan fluorescence and far-UV CD. The experiments were performed in the presence of POPC:POPG:chol 7:2:1 SUV. Data for P233 and P414 as a function of lipid concentration are displayed in magenta and blue, respectively. Membrane partitioning followed by 1D ^1^H NMR is represented on a linear scale (A) and on a logarithmic scale (B). The ratio of fluorescence intensity (320 nm/370 nm) followed by tryptophan fluorescence (C) and circular dichroism in the far-UV region monitored by far-UV CD (D) are shown. The signal loss in (A) and (B) of the tryptophan indol proton (open circles) followed by NMR is similar to that of the amide envelope (closed circles).

Interestingly, we noticed that, while in many cases (*e*.*g*., apo-Mb, ALO or peptide P233) no residual NMR signal is observed at saturating lipid concentrations (*i*.*e*., after reaching a plateau), a significant NMR signal remains for the T domain at the highest lipid concentrations tested. This signal might be due to either a fraction of protein unable to bind to membranes and remaining in solution or, alternatively, it might arise from fully membrane-attached polypeptides containing flexible regions not directly bound to the lipids and floating above the membrane (disordered regions or ordered regions linked to the membrane-bound region(s) by a flexible linker, see figure 6). To discriminate between these two possibilities, we performed NMR self-diffusion experiments as diffusion is expected to be different for proteins in solution or bound to the lipid vesicles.

**Figure 6.**
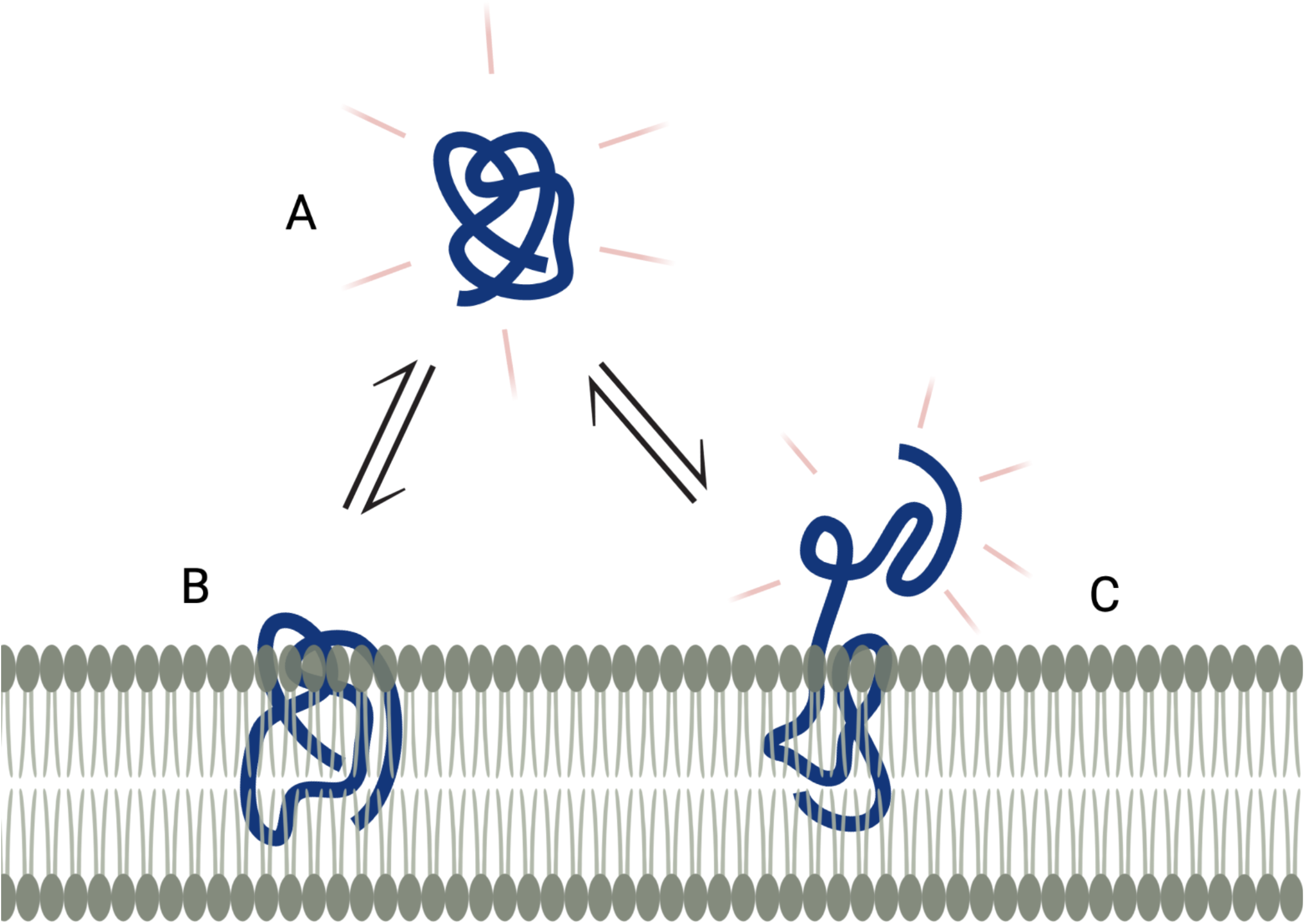
Schematic model of three potential conformational states of an amphitropic protein exposed to membranes. The experimental results of the B2LiVe method report on the fraction of amide signal (red traces) that is lost upon titration of the protein (black line) titration by membranes. The total loss of the NMR signal upon protein titration by lipid membranes indicates that all proteins populate the B state at the expense of the A state. If the NMR signal does not fully disappear and reaches an intermediate plateau, this indicates that a fraction of the proteins remains in solution. The remaining amide signal may arise either from a fraction of the population of proteins remaining in solution (A state) or from fully membrane-bound proteins that contain specific regions remaining in solution (C state). The DOSY experiment allows to discriminate between these possibilities.

Diffusion experiments were performed with the T domain (15 *μ*M) alone or in the presence of a 1000-fold excess of lipids in LUV at pH 4.5. Under these conditions, in amide-selective 1D spectra, we observed that in the presence of LUV, the signal was ∼40% (39 ± 3 %) that of the protein without lipid vesicles (Figure 7A). We then submitted the samples to amide-selective diffusion experiments. In diffusion experiments in which all delays (diffusion delay, gradient pulses) are kept constant and only the gradient strength is varied, the signals show a gaussian decay that depends on the diffusion coefficient and the applied gradient. As can be observed in Figures 7B and 7C, the relative decay on intensity of the spectra at high gradient strength is much higher for the T domain alone that in the presence of LUV. This can also be visualized on the diffusion curves (Figure 7D), which show the signal intensity as a function of the gradient strength. These data indicate that the diffusion coefficient of the species giving rise to the residual signal at high lipid concentration are bound to lipid vesicles. To rule out the possibility that the slower diffusion of the T domain at high lipid concentration could be due to a higher viscosity resulting from the presence of LUV rather than a consequence of membrane binding, we used a buffer signal (Hepes resonance) as a control and showed that Hepes displays very similar diffusion decays with or without lipid vesicles in samples containing the T domain (Figure 7E). Hence, the residual signal pertains to membrane-bound protein, in agreement with previous results ^10^. In summary, in addition to quantifying membrane partitioning, the B2LiVe method coupled to diffusion-ordered spectroscopy (DOSY) can pinpoint toward the presence, within a membrane-bound protein, of polypeptide segments that are not imbedded into membrane, but remain in the aqueous phase.

**Figure 7.**
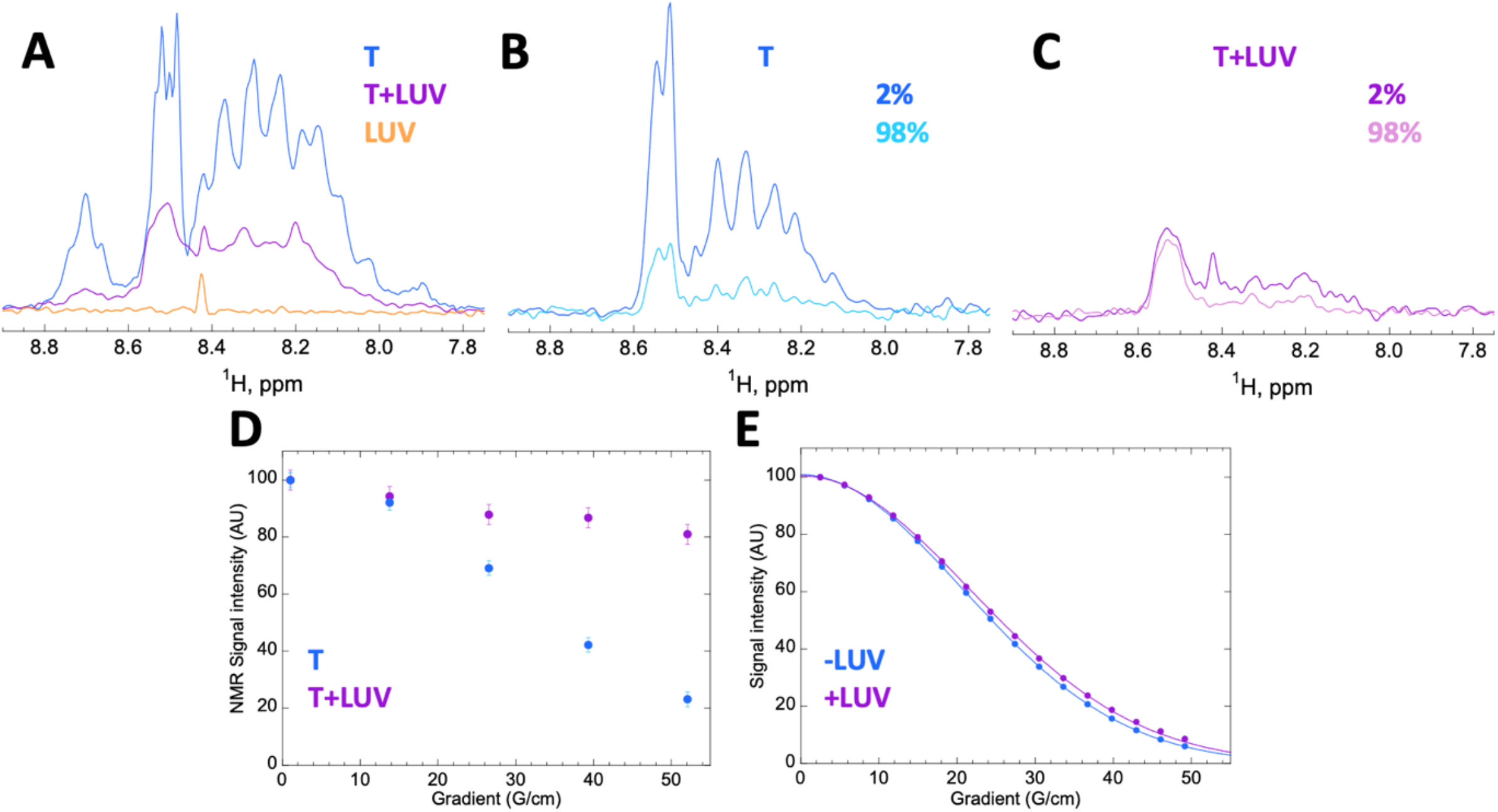
Self-diffusion of the T domain in the absence and presence of LUV monitored by 1D ^1^H amide-selective diffusion NMR experiments. A. 1D ^1^H amide-selective spectra of domain T (15 *μ*M) at pH 4.5 and 25°C in the absence (blue) or presence (violet) of LUV (15 mM concentration), and of LUV (POPC:POPG 9:1) without protein (orange). The envelope integral decays to 39 ± 3 % in the presence of LUV. (B to D) diffusion experiments recorded with a 140 ms diffusion delay. 1D Spectra at 2% and 98% of the maximum gradient available (53.5 G/cm) of T alone (B) or in the presence of LUV (C) displayed on the same scale. (D) Diffusion rate-dependent decay of the intensity (integral) of the signal at 8.55 ppm of T with (violet) or without (blue) LUV. The much lower decrease in relative intensity of the signal in the presence of LUV indicates a slower diffusion rate. (E) diffusion rate-dependent decays of the Hepes signal (3.86 ppm) in samples (used in A to D) with (violet) or without (blue) lipid vesicles. Experiments in (E) were recorded with a 60 ms diffusion delay; data in (E) were fitted to a gaussian decay of the intensity (I) from an Io value (without gradients) as a function of the gradient strength G and an apparent diffusion coefficient d (*I* = *Ioe*^−*dG*2^). The similarity of the diffusion decays in (E) indicates that the viscosity of the sample is not significantly increased by the LUV and is not at the origin of the slower diffusion of T in the presence of lipid vesicles shown in B, C and D. The sharp signal at 8.43 ppm arises from a low molecular weight trace impurity.

## Discussion

We have shown that 1D ^1^H NMR with selective excitation of the amide region is a simple and robust approach to quantify membrane partitioning of unlabeled peptides or proteins. Selective excitation of the amide region allows one to efficiently filter-out the proton signals of water (111 M), buffers (tenths of millimolar range), lipid vesicles (from 0 to 10 mM in this study) as well as most of the impurities that are usually observed at lower frequencies, without the need of ^15^N-labeling the proteins. In addition, the fast relaxation of the amide protons in the HET-SOFAST experiment ^1^ warrants the use of short inter-scan repetition delays leading to high sensitivity. With high-field spectrometers and high-sensitivity, cryogenically cooled probes, data acquisition on low protein-concentration samples can indeed be very short. In the experiments presented here, run on an 800 MHz spectrometer equipped with a cold probe, one data point was recorded with 2048 scans in ∼12 minutes for 3 *μ*M protein in 180 *μ*L samples.

The B2LiVe method requires low concentrations and low amounts of unlabeled proteins, is independent of the protein/peptide composition and can be applied to a large range of protein sizes, which can go well-above the range of the protein sizes shown in this study (> 54 kDa, unpublished data). Importantly, the capacity to perform the experiments at low protein concentrations and high lipid/protein ratios, ensures the conditions of protein ‘infinite’ dilution required to determine the thermodynamic partitioning constants (see Equation 2). On the other hand, given that the determination of the membrane-bound fraction is based on the disappearance of the solution protein signals, which is due to the large size of the lipid vesicles, the method is independent of the lipid bilayer particle size (SUV or LUV) and is not negatively affected by high lipid concentration, which can cause light scattering problems in optical techniques like fluorescence and CD. This tolerance to particle size and high concentration allows the technique to quantify the solution-to-membrane partitioning for systems with low affinity.

Furthermore, as we showed here for the T domain, and in contrast to other techniques, NMR can directly indicate if certain regions from a membrane-bound protein remain in the aqueous phase. In its molten globule state at acidic pH, the T domain is able to penetrate membranes on a pH dependent manner ^26^. Here, we showed that at pH 4.5, *ca*. 40% of the membrane-bound protein residues (39 ± 3 % residual amide signal) are not directly attached to the membranes and remain flexible enough in solution so that their NMR signals can be observed. Previous works on the T domain, as well as with peptides encompassing its four N-terminal helices (TH1 to TH4), indicate that under the conditions used in this study (pH 4.5, POPG:POPC 9:1 LUV), the N-terminal region does not interact with membranes ^10,11^; membrane insertion of the N-terminal helices of T is only observed for more acidic pH (pH ≤ 4) ^26,11^. Remarkably, this N-terminal amphiphilic region corresponds to ∼40 % of the residues of the T domain, in close agreement with our NMR results (39 ± 3 % residual amide signal).

In conclusion, the proposed NMR-based B2LiVe method should be valuable to quantify the affinity of unlabeled peptides and proteins for lipid bilayers in the context of structural biology and biophysical studies.

### Limitations of Study

The B2LiVe method may be difficult to apply for proteins or peptides that have a strong propensity to aggregate. However, for such peptides/proteins, other classical approaches such as SPR, reflectometry-based or centrifugation-based partitioning assays will also be severely impeded and only membrane flotation assays might potentially be applicable to demonstrate their membrane binding capacity. However, it should be stressed that because B2LiVe can be carried out at high lipid:polypeptide molar ratio (*i*.*e*., in the low micromolar range for proteins and in the tens of millimolar range for lipids), the aggregation-propensity of most proteins should be rather limited in such conditions. Also, if there is intermediate exchange, affinity could be overestimated but this effect should be easily detectable by visual inspection of the NMR spectra. In this latter case, the B2LiVe method will still provide a good estimation of the affinity range (i.e. order of magnitude of K_D_) that should be useful for performing most comparative studies (e.g. effects of mutations, different lipid compositions, …).

## Acknowledgements

M.S. was supported by the Pasteur -Paris University (PPU) International PhD Program. N.C. was supported by Institut Pasteur (DARRI-Emergence S-PI15006-12B). The 800 MHz NMR spectrometer of the Institut Pasteur was partially funded by the Région Ile de France (SESAME 2014 NMRCHR grant no 4014526). We also acknowledge funding from the Agence Nationale de la Recherche (ANR 21-CE11-0014-01-TransCyaA), the CNRS (UMR 3528) and the Institut Pasteur (PTR 166-19, DARRI-Emergence S-PI15006-12B, PPUIP program). We thank W. J. Tang (Chicago University, USA) for the kind gift of purified ALO and Bruno Baron and the PFBMI platform at Institut Pasteur for assistance in the performance of the far-UV CD experiments. The funders have no role in study design, data collection and analysis, decision to publish, or preparation of the manuscript.

## Author contributions

Conceptualization, J.I.G. and A.C., Methodology, J.I.G. and A.C., Software, B.V. and J.I.G., Validation, B.V., D.L., J.I.G. and A.C., Formal Analysis, M.S., N.C., B.V., J.I.G. and A.C., Investigation, M.S., N.C., B.V. and J.I.G., Resources, D.L., J.I.G. and A.C., Writing – Original Draft, M.S., D.L., J.I.G. and A.C., Visualization, M.S., N.C., J.I.G. and A.C., Supervision, J.I.G. and A.C., Project Administration, J.I.G. and A.C., Funding Acquisition, J.I.G. and A.C.

