## Supplemental Tables for "B2LiVe, a label-free 1D-NMR method to quantify the binding of amphitropic peptides or proteins to membrane vesicles"

*Corresponding authors

Supplemental Table S1

**Table S1: Properties of the proteins used in this study**

| Protein | Molecular mass, * kDa | Isoelectric point | Molar extinction coefficient, * $M^{-1}.cm^{-1}$ |
|---|---|---|---|
| BLA | 14.2 | 4.5 | 20100 |
| apo-CaM | 16.7 | 4.1 | 2980 |
| apo-Mb | 16.9 | 7.4 | 13980 |
| T | 22.0 | 5.5 | 17780 |
| ALO | 54.0 | 6.3 | 62800 |
| BSA | 66.4 | 4.7 | 43824 |

* Obtained with Protparam (https://web.expasy.org/protparam) and the product information sheets provided by Sigma Aldrich.

Abbreviations: bovine α-lactalbumin (BLA), apo-calmodulin (apo-CaM), apo-myoglobin (apo-Mb), diphtheria toxin translocation domain (T), anthrolysin O (ALO), bovine serum albumin (BSA).

Supplemental Table S2

**Table S1: Properties of the peptides used in this study**

| Peptide name | Number of residues | Molecular mass, kDa* | Isoelectric point ** | Charge at neutral pH ** | Molar extinction coefficient, ** $M^{-1}.cm^{-1}$ | Maximum hydrophobic moment, *** $\mu H_{max}$ | Hydrophobicity <H> corresponding to $\mu H_{max}$ *** |
|---|---|---|---|---|---|---|---|
| P233 | 22 | 2.5 | 10.7 | + 2 | 5500 | 0.384 | 0.223 |
| P414 | 27 | 2.9 | 4.25 | - 4 | 5500 | 0.224 | 0.324 |

* Determined by MALDI-TOF mass-spectrometry
** Obtained with Protparam (https://web.expasy.org/protparam)
*** The values for $\mu H_{max}$ and of the corresponding <H> were obtained via Heliquest (https://heliquest.ipmc.cnrs.fr/) using a window of 18 amino acids.

Supplemental Table S3

**Table S3: Protein-partitioning thermodynamic data**

| Protein | Methods used | Membrane partitioning coefficient, $K_x$ | Dissociation constant $K_D$, μM | Hill coefficient, nH | Free energy $-\Delta G_{Kx}$, kcal/mol |
|---|---|---|---|---|---|
| apo-Mb | NMR | 170000 | 333 | 2 | 7.1 |
| | fluorescence | 77000 | 608 | 2 | 6.7 |
| T | NMR | 240000 | 226 | 3 | 7.4 |
| | fluorescence | 250000 | 218 | 2 | 7.4 |
| ALO | NMR | 800000 | 67 | 2 | 8.1 |
| | fluorescence | 640000 | 86 | 3 | 7.9 |

Thermodynamic data determined from experimental data obtained by NMR and tryptophan fluorescence. The standard deviation values are < 10%. Abbreviations: apo-myoglobin (apo-Mb), diphtheria toxin translocation domain (T), anthrolysin O (ALO).

Supplemental Table S4

**Table S4: Peptide P233 partitioning thermodynamic data**

| Methods used | Membrane partitioning coefficient, $K_x$ | Dissociation constant $K_D$, µM | Free energy $-\Delta G_{Kx}$ , kcal/mol | Hill coefficient, nH |
|---|---|---|---|---|
| **NMR** | 52000 | 1000 | 6.4 | 3 |
| **Fluorescence** | 26000 (13000) | 2100 (4100) | 6.0 (5.6) | 2 (18) |
| **CD** | 25000 | 2200 | 6.0 | 3 |

Thermodynamic data were determined from experimental data obtained by NMR, tryptophan fluorescence and far-UV CD. The fluorescence experimental data set was fitted assuming two transitions. The thermodynamic values describing the second transition, related to peptide reorganization in the membrane, are reported in brackets.
